# Kasugamycin inhibits melanoma lung metastasis and regulates CHI3L1-driven M2-like tumor-associated macrophage differentiation

**DOI:** 10.1101/2025.05.26.654629

**Authors:** Takayuki Sadanaga, Han-Seok Jeong, Roberto Cortez, Joyce H Lee, Suchitra Kamle, Bing Ma, Sung Jae Shin, Jack A. Elias, Chun Geun Lee

## Abstract

CHI3L1, a chitinase-like protein, is a potent immune modulator involved in various diseases, including lung cancer. While recent studies have demonstrated that kasugamycin (KSM) is a pan-chitinase inhibitor with strong anti-fibrotic activity, its effects on specific chitinase-like proteins remain undefined. This study shows that KSM effectively abrogates CHI3L1-stimulated cellular signaling and bioactivities. In a B16/F10 melanoma lung metastasis model, where CHI3L1 plays a critical role, KSM treatment significantly reduced melanoma lung metastasis dose-dependently. The anti-tumor effect of KSM was found to be CHI3L1-specific, as CHI3L1 overexpression enhanced melanoma lung colony formation, which was effectively blocked by KSM. In melanoma-challenged lungs, KSM treatment significantly reduced the elevation of M2 macrophages expressing CD206, CD163, and PD-L1. In studies using human monocytic THP-1 cells, CHI3L1 promoted M2 macrophage differentiation, which KSM significantly suppressed. Bulk RNA sequencing of differentiated macrophages revealed that CHI3L1 highly induced the expression of epidermal growth factor receptor (EGFR), and this induction was counter-regulated by KSM, and CHI3L1-driven M2 macrophage activation was reduced with EGFR blocker treatment. These findings reveal a novel anti-tumor mechanism of KSM, which inhibits M2-like tumor-associated macrophage differentiation, potentially through the CHI3L1-EGFR axis.

## Introduction

Lung cancer is the leading cause of cancer-related mortality worldwide for both men and women, accounting for approximately 1 in 5 cancer deaths in the United States (1). Non-small cell lung cancer (NSCLC) comprises 80–85% of all lung cancer cases, with over half of patients being diagnosed at an advanced stage of the disease (2). Additionally, the lung is the second most frequent site for metastatic spread, with 20–54% of malignant tumors originating elsewhere in the body leading to pulmonary metastases (3). Despite significant advances in lung cancer management, including the use of immune checkpoint inhibitors, the 5-year survival rate for advanced-stage cases remains under 10%. Disease progression during or after treatment continues to be the primary cause of mortality (3). Therefore, developing effective therapeutic strategies to block disease progression and improve patient outcomes remains a critical unmet need.

The glycosyl hydrolase family 18 (GH18) includes true chitinases, such as Chitinase 1 (CHIT1), and several chitinase-like proteins (CLPs) that lack enzymatic activity. Among these, Chitinase 3-like 1 (CHI3L1, also known as YKL-40) is a prototypic CLP that is overexpressed in the serum and tissues of lung cancer patients. Elevated serum levels of CHI3L1 have been identified as an independent prognostic biomarker for poor survival in lung cancer patients (4). CHI3L1 has been shown to drive a pro-tumoral microenvironment through modulation of innate and adaptive immune responses in various cancers, including lung cancer (5–11). Recent studies from our laboratory revealed that CHI3L1 stimulates the expression of immune checkpoint molecules and their receptors, including PD-1/PD-L1 and CTLA-4, in macrophages and T cells in the lung, contributing to enhanced melanoma lung metastases (10, 11). Among immune cells, macrophages, particularly M2-like tumor-associated macrophages (TAMs), are a major component of the tumor microenvironment. These cells, which highly express CHI3L1, play a crucial role in tumor progression, and CHI3L1’s involvement in promoting M2 macrophage differentiation has been proposed (5, 12). However, the direct role and mechanism of CHI3L1 in specific macrophage differentiation, driving its pro-tumoral effects and potential therapeutic interventions, have not been sufficiently explored.

Kasugamycin (KSM) is a naturally occurring aminoglycoside antibiotic characterized by two sugar moieties and low animal toxicity (13). Recently, KSM was identified as a specific inhibitor of chitinase, demonstrating strong anti-fibrotic activity by blocking CHIT1-stimulated TGF-β signaling in the lung (14, 15). KSM has also been shown to enhance host resistance to viral infections by upregulating interferon-stimulated genes (ISGs) (16). Further studies have revealed KSM’s broad anti-chitinase activity across various chitinases from humans and other organisms (17), highlighting its potential use as a chitinase inhibitor. However, its activity against CHI3L1 and other GH18 family members that share structural and functional similarities has not been determined.

In this study, we demonstrate that KSM significantly suppresses melanoma lung metastases. Our findings reveal that CHI3L1 drives M2 macrophage differentiation, a process counter-regulated by KSM treatment. Specifically, CHI3L1 increased the number of macrophages expressing CD206, CD163, and PD-L1, hallmarks of M2-like TAMs in melanoma lung metastases. This effect was more pronounced in CHI3L1-overexpressing transgenic mice compared to wild-type controls. We further identified that CHI3L1 stimulates M2 macrophage differentiation, which was significantly inhibited by KSM, potentially through regulation of EGFR expression. Collectively, these studies highlight a novel anti-tumor mechanism of KSM, which inhibits CHI3L1-mediated protumor activity in M2 macrophage differentiation, thereby suppressing melanoma lung metastases.

## Results

### Kasugamycin inhibits melanoma lung metastasis

To evaluate the effects of KSM on pulmonary melanoma metastasis, wild-type (WT) mice were injected via the tail vein with either B16-F10 melanoma cells (B16) or vehicle control (PBS), as illustrated in Fig. 1A. Fourteen days post-challenge, metastatic melanoma colonies (pleural colonies) were prominent in mouse lungs and were counted and compared between PBS-treated mice and those treated with varying doses of KSM (Fig. 1B). These studies revealed that KSM significantly attenuated melanoma lung metastasis in a dose-dependent manner. Hematoxylin and eosin (H&E) staining further demonstrated reduced parenchymal tumor growth in the lungs of KSM-treated mice compared to PBS-treated controls (Fig. 1C). Additionally, the degree of inhibition in melanoma lung metastasis correlated positively with the duration of KSM treatment (Supplemental Fig. S1).

**Figure 1.**
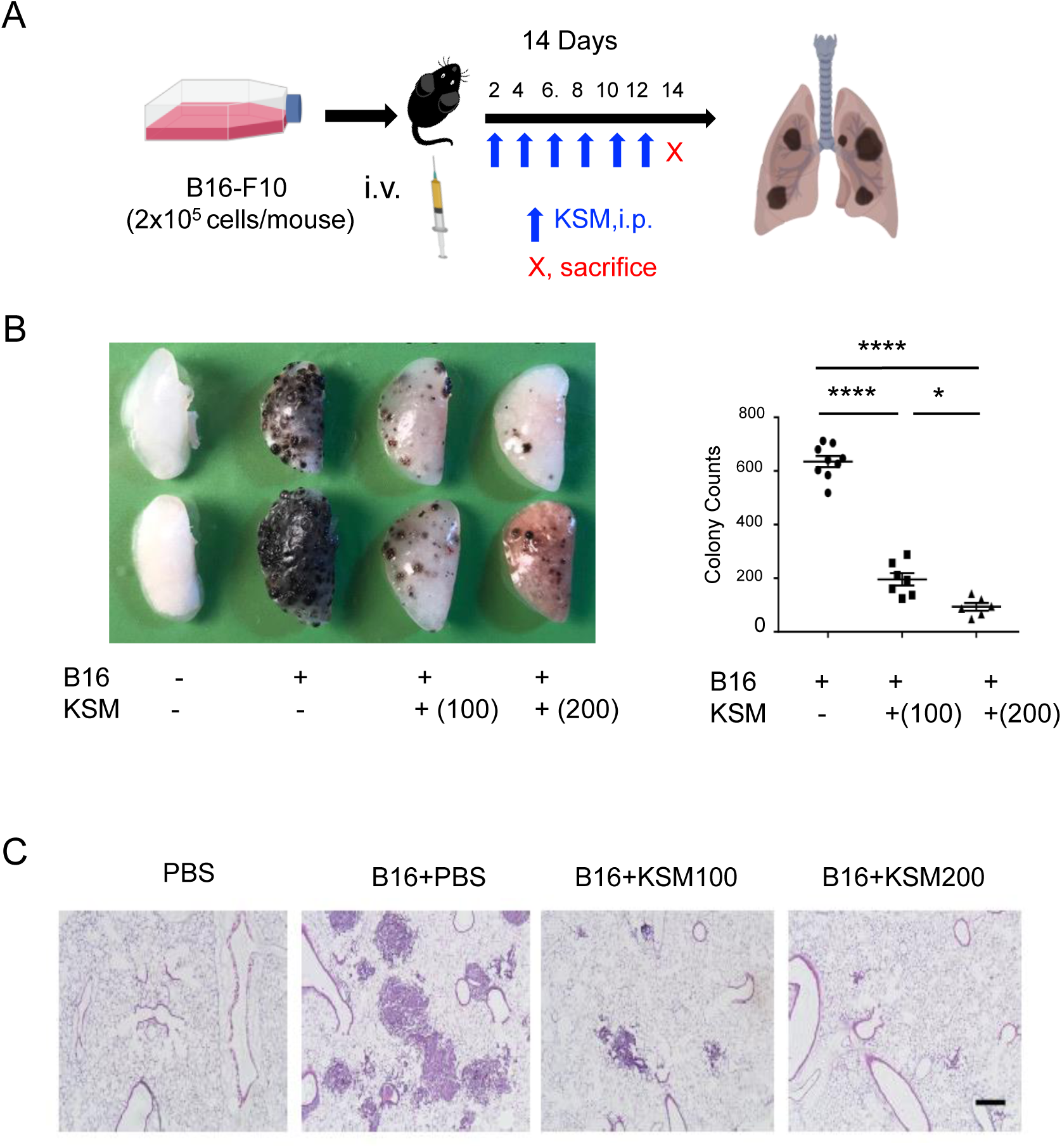
Kasugamycin inhibits melanoma lung metastasis. (A) Schematic illustration of melanoma lung metastasis evaluation. (B) Dose-dependent kasugamycin (KSM) inhibition of melanoma lung metastasis. B16-F10 (B16) melanoma cells were delivered to the mice through tail vein injection with and without KSM treatment (100 mg/kg and 200 mg/kg mouse, i.p.). Representative lung photos with melanoma lung metastasis (left panel) and their pleural colony counts (right panel). (C) Representative H&E stains of B16-F10 (B16) melanoma cell-challenged lungs with and without KSM treatment. The values in panel B for the pleural colony count are the mean ± SEM. *p < 0.05, ****p < 0.0001 (One-Way ANOVA, multiple comparisons). Scale bar in panel C = 250 µm

### Kasugamycin inhibits CHI3L1-mediated signaling and anti-apoptotic responses of lung macrophages

Given that KSM is a potent inhibitor of CHIT1 and other enzymatically active chitinases (17, 18), we hypothesized that KSM may similarly influence the signaling and biological activity of CHI3L1. To test this, we first investigated the impact of KSM on well-characterized CHI3L1 signaling pathways (19). As shown in Fig. 2A, recombinant CHI3L1 (rCHI3L1) enhanced the phosphorylation of Erk and Akt in AMJ2-C11 mouse lung macrophages. These CHI3L1-stimulated activations were effectively abrogated by KSM treatment, similar to anti-CHI3L1 antibody (FRG) treatment. CHI3L1 is also known for its anti-apoptotic effects, particularly in reducing oxidative stress-induced apoptosis (19). To evaluate KSM’s impact, macrophages were treated with hydrogen peroxide (H_2_O_2_) to induce oxidative stress, and apoptosis was assessed by fluorescence-activated cell sorting (FACS) analysis. CHI3L1 treatment significantly decreased the number of Annexin V (+) apoptotic lung macrophages. However, this anti-apoptotic effect was significantly reduced upon KSM treatment to levels comparable to anti-CHI3L1 antibody (FRG) treatment (Fig. 2B), suggesting that KSM effectively blocks CHI3L1-mediated anti-apoptotic signaling in macrophages. Together, these findings demonstrate that KSM effectively inhibits CHI3L1-mediated signaling and cellular responses in macrophages.

**Figure 2.**
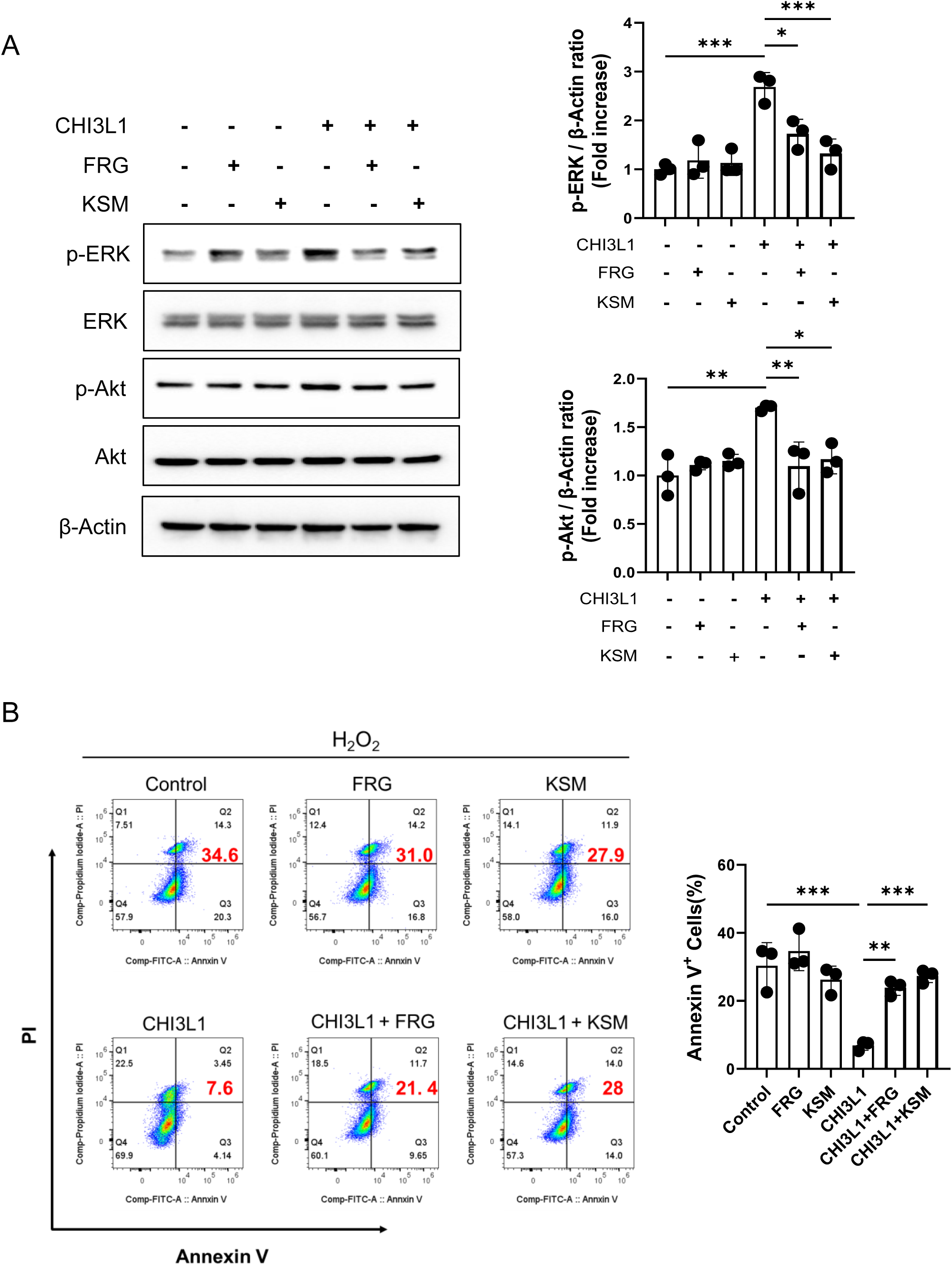
KSM inhibits CHI3L1-stimulated Erk and Akt activation and antiapoptotic effect of CHI3L1 in macrophages. (A) AMJ2-C11 mouse lung macrophages were stimulated with recombinant (r) CHI3L1 overnight, and the activation of Erk and Akt was detected by Western blot evaluations. pErk, phosphorylated Erk; T-Erk, total Erk; pAkt, phosphorylated Akt, T-Akt, total Akt. Right panel, densitometric quantitation on the blots of pErk and pAkt. (B) After exposure of AMJ2-C11 cells with H_2_O_2_ (1 mM, 24 hours) with and without rCHI3L1 and KSM treatment, cellular apoptosis (% of Annexin V+ cells) was measured by FACS evaluations. Panels A and B are representatives of 3 independent experiments. Right panel, the values of Annexin V (+) cells (%) are the mean ± SEM. **p < 0.01, ***p < 0.001 (One-Way ANOVA, multiple comparisons).

### KSM inhibits M2-like TAM accumulation in lungs with melanoma metastasis

To investigate the roles of CHI3L1 and KSM in macrophage differentiation during melanoma lung metastasis, we characterized macrophages accumulated in the lungs of WT and CHI3L1 transgenic mice following B16-F10 challenge and KSM treatment. Immunohistochemistry and FACS analyses revealed increased numbers of macrophages expressing CD206, CD163, and PD-L1 (markers of M2-like TAMs, (20, 21)) in the lungs of B16-challenged mice (Fig. 3). Notably, KSM treatment significantly reduced CD206(+)/CD163(+) macrophages in peritumoral regions, indicating that KSM inhibits M2-like TAM accumulation in the lung (Fig. 3A). PD-L1 expression was similarly elevated in CD206(+) and CD163(+) macrophages within the lungs of melanoma-challenged mice. However, it was markedly reduced following KSM treatment (Fig. 3B).

**Figure 3.**
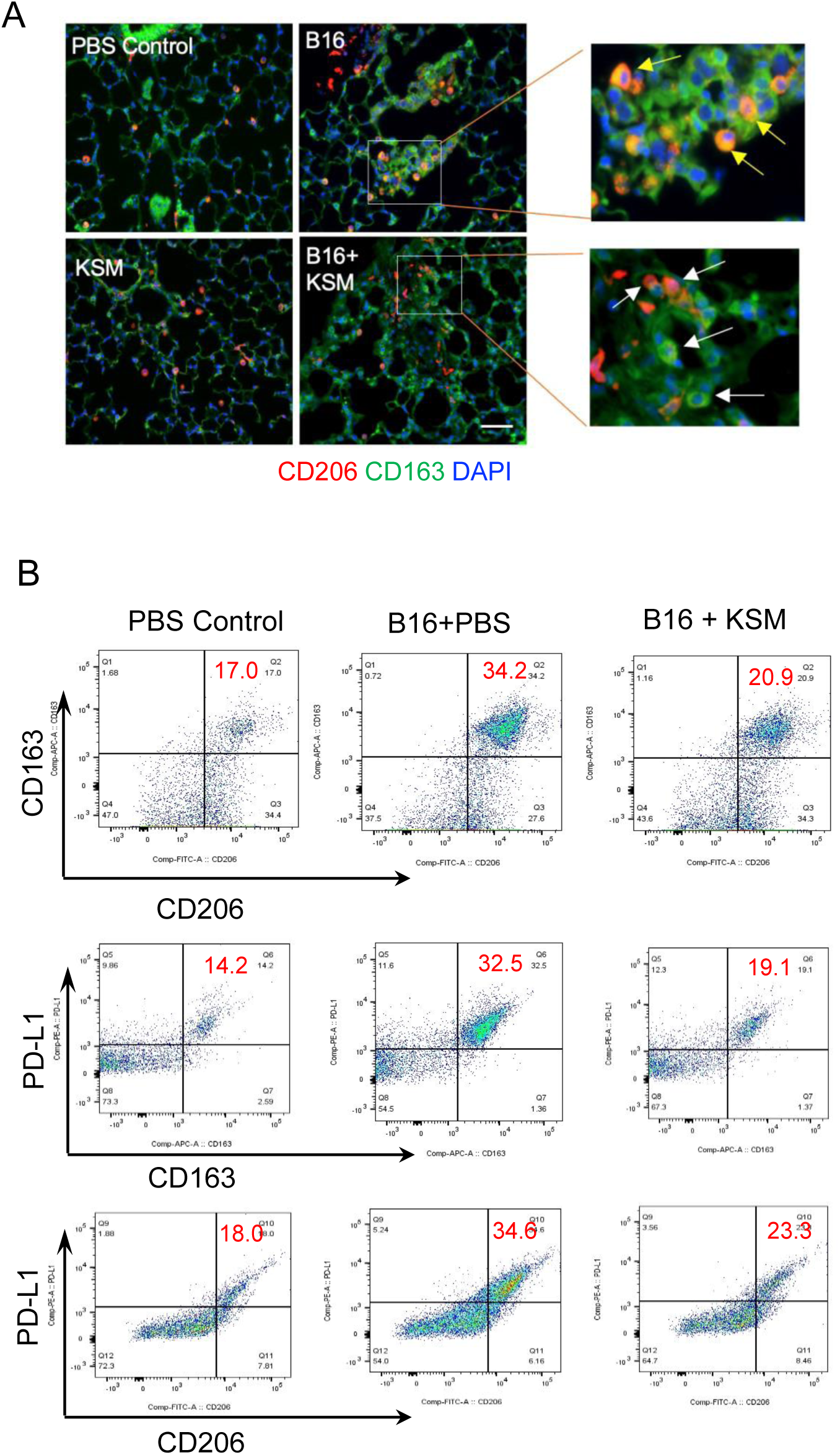
KSM inhibits M2-like TAMs accumulation in lungs of melanoma metastasis. (A) Co-immunostaining of CD206 (red) and CD163 (green) in lungs challenged with B16-F10 (B16) melanoma cells with vehicle (PBS) or KSM treatment (100 mg/kg/mouse). Arrows indicate CD206/CD163 double (+) cells (yellow) or CD206 or CD163 single positive cells (white). (B) Representative FACS evaluations on CD206(+)/CD163(+), CD206(+)/ PD-L1(+), and CD163(+)/PD-L1(+) macrophages in lungs with PBS or melanoma cell challenge and KSM treatment. Scale bar in Panels A = 25 µm

### CHI3L1 enhances the melanoma lung metastasis and M2 macrophage accumulation in the lung

To see the impact of CHI3L1 in melanoma lung metastasis and tumor-associated macrophage activation, we evaluated the WT and CHI3L1 Tg mice after low-dose B16-F10 melanoma cell challenge (0.5×10^5^ cells/mouse). The number of melanoma lung colonies was significantly increased in CHI3L1 Tg mice compared to WT mice, and the increases were abrogated by KSM treatment (Fig. 4, A and B). Immunohistochemical evaluations further revealed a significant accumulation of CD206(+)/CD163(+) and CD206(+)/PD-L1(+) cells in the lungs of CHI3L1 Tg mice, and these increases were abrogated by KSM treatment (Fig. 4, C and D). These findings suggest that CHI3L1 promotes M2 or M2-like TAM accumulation in melanoma lung metastases and that KSM effectively counteracts this process.

**Figure 4.**
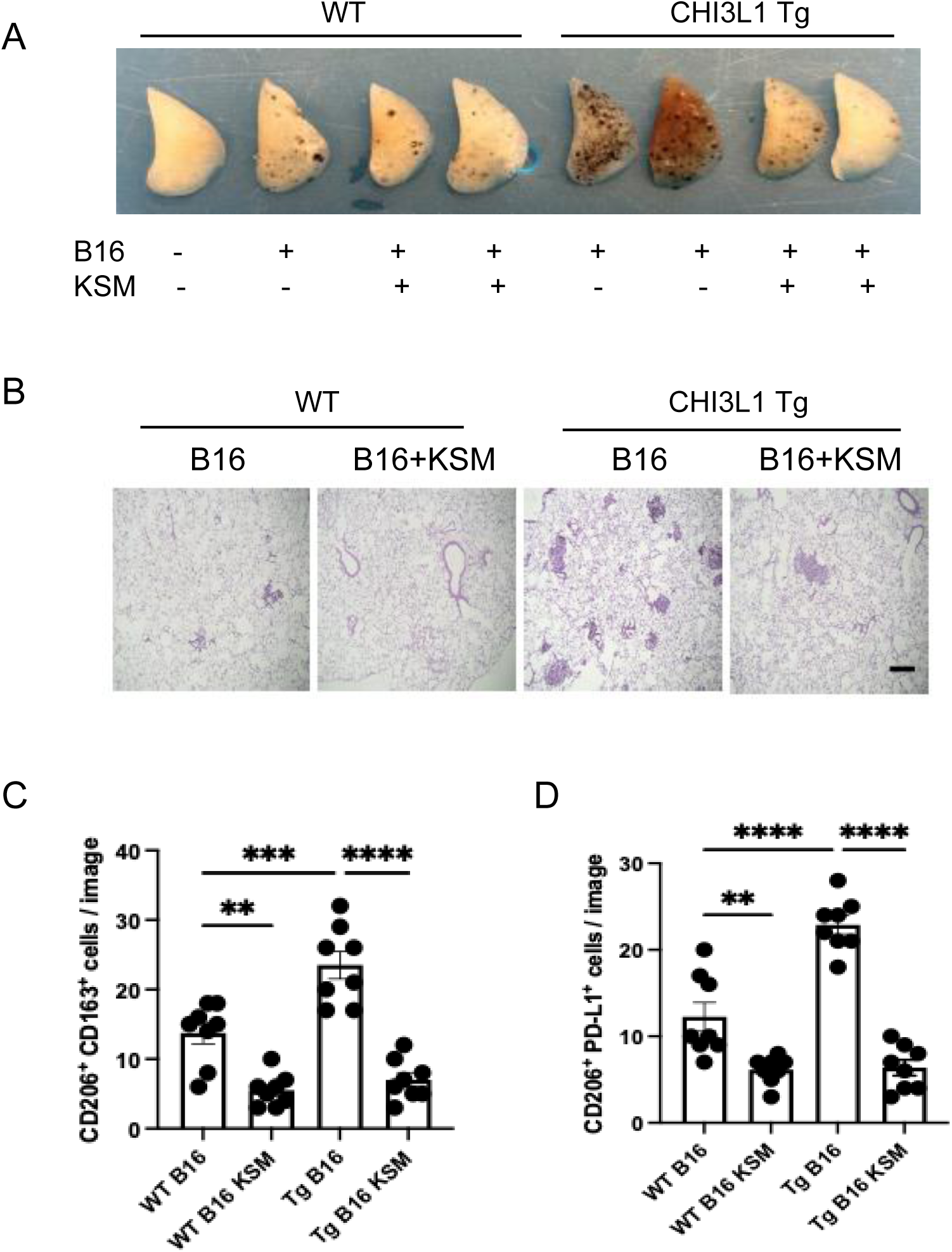
KSM inhibits CHI3L1-induced M2 macrophage accumulation in melanoma lung metastasis. (A). Representative photographs of WT and CHI3L1 transgenic lungs with and without KSM treatment after B16-F10 melanoma cell challenge. (B) Representative H&E histology of WT and Transgenic lungs with and without KSM treatment after B16-F10 melanoma cell challenge (x4 original magnification). (C-D) The number of CD206(+)/CD163(+) and CD206(+)/PD-L1(+) macrophages detected by immunohistochemical staining in lungs of WT and CHI3L1 Tg mice with and without KSM treatment. The number of macrophages co-expressing CD206 and CD163 or PD-L1 was counted using microscopic images under 20x magnification (n = 8 each). The values in panels C and D are the mean ± SEM. *p < 0.05, **p < 0.01 ***p < 0.001 ****p < 0.0001 (One-Way ANOVA, multiple comparisons).

### KSM inhibits CHI3L1-stimulated M2 macrophage differentiation *in vitro*

To further investigate CHI3L1’s capacity to promote M2 or M2-like TAM differentiation, we conducted in vitro studies using THP-1 human monocytic cells. After 24 hours stimulation with Phorbol 12-myristate 13-acetate (PMA) followed by additional 24 hours resting period, THP-1 cells were exposed to rCHI3L1 for 48 hours in the presence and absence of tumor-conditioned media (TCM; supernatant from A549 cells) (Fig. 5A). The expression of M2 or M2-like TAM activation markers, including CD206, CD163, CD274 (PD-L1), CX3CR1 and CD36 was assessed by real-time qRT-PCR evaluations. These experiments revealed that rCHI3L1 upregulated expression of typical markers of M2 or M2-like TAM differentiation with no significant effect on the expression of NOS2, a marker of M1 macrophage differentiation (Fig. 5, B, and C and Supplemental Fig. 2). Importantly, KSM treatment significantly suppressed CHI3L1-stimulated M2 activation marker expressions regardless of TCM treatment (Fig. 5, B and C). Immunoblot evaluation further confirmed that CHI3L1 induces CD206, CD163, and PD-L1 protein expression, which is strongly inhibited by KSM treatment (Fig. 5D).

**Figure 5.**
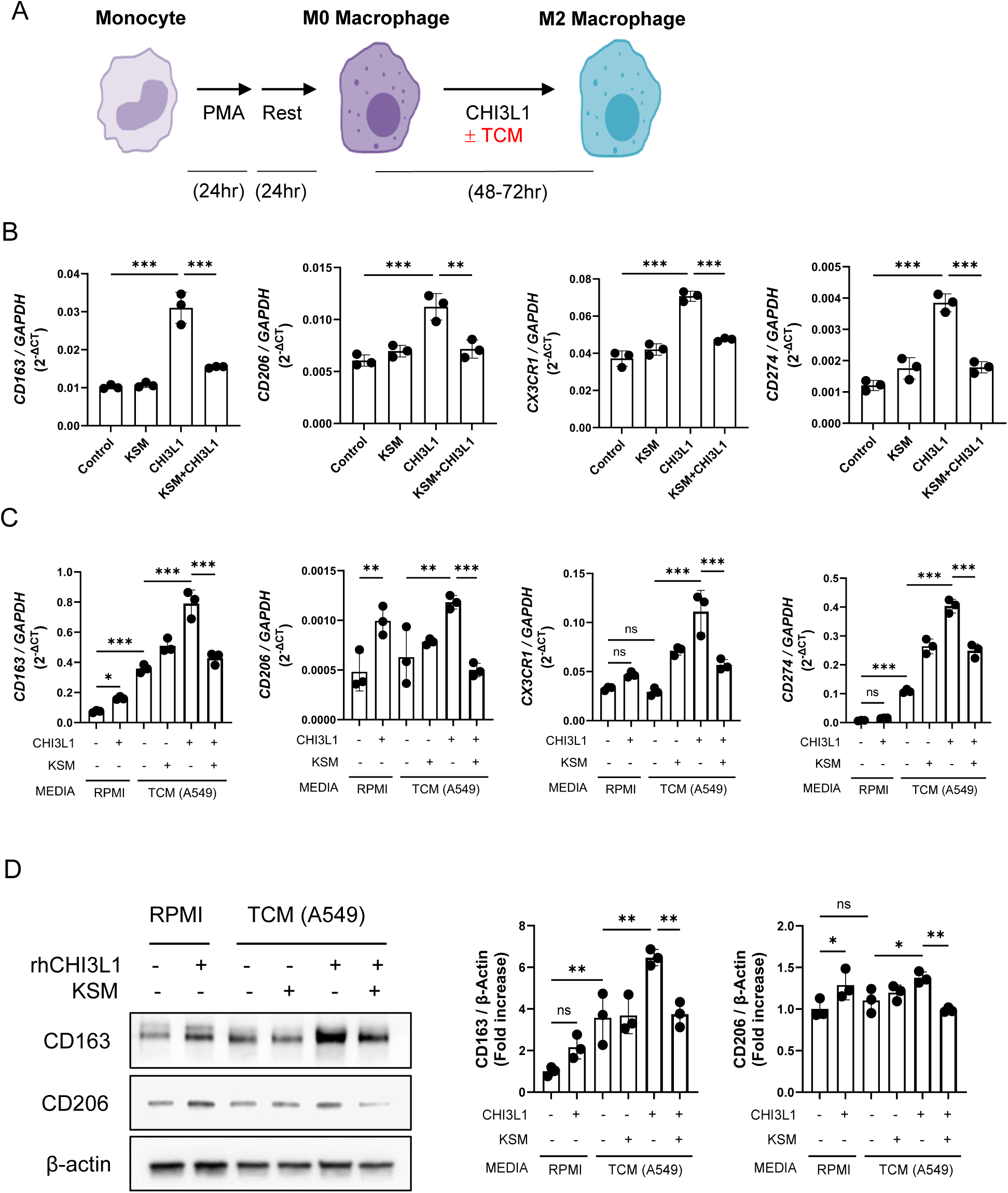
KSM inhibits CHI3L1-stimulated differentiation of monocytic macrophages to M2 or M2-like TAM macrophages. (A.) Schematic illustration of macrophage differentiation using human THP-1 cells. (B) Real-time PT-PCR evaluations on the M2 macrophage activation markers in the macrophages with and without CHI3L1 (500 ng/mL) and KSM (250 ng/mL) treatment. (C) Real-time qRT-PCR evaluations on the M2 macrophage activation markers in the macrophages cultured in the presence and absence of tumor cell conditioned medium (TCM; A549 cell supernatant), together with CHI3L1 and KSM treatment. (D) Representatives of immunoblot evaluations on the expression of CD206/CD163 in differentiated macrophages stimulated in the presence and absence of TCM, together with CHI3L1 and or KSM treatment. Right panel, densitometric quantitation on the blots of CD163 and CD206. The values in panels B, C, and D are mean ± SEM. *p < 0.05, **p < 0.01 ***p < 0.001 (One-Way ANOVA, multiple comparisons).

### Identification of KSM-regulated genes in CHI3L1-driven M2 macrophage

To elucidate the molecular mechanisms underlying KSM’s anti-tumor activity, we performed mRNA expression profiling using bulk RNA sequencing on differentiated macrophages in the presence of CHI3L1 alone or in combination with KSM. Differential gene expression analysis identified 519 upregulated and 494 downregulated genes (≥ 2-fold change) in rCHI3L1-treated macrophages compared to PBS controls. As illustrated by volcano plots (Fig. 6A), CHI3L1 stimulated the expression of multiple genes including metalloproteinases (MMP-8, MMP-9), growth factor (EGFR), chemokine (CXCL13), and other signaling molecules, which are associated with pathways involved in anti-apoptosis, proliferation, adhesion, and immune modulation (Supplemental Table S1 and Supplemental Fig. S3). On the other hand, KSM treatment significantly increased the expression of genes associated with protein translation and mitochondrial function, including ribosomal genes such as RPL41P1, RPL35, PVALB, and ROMO1 (Supplemental Table S2 and Supplemental Fig. S4). Interestingly, several genes associated with protumor activities, such as EGFR, FP671120.4, ITGB8, TNC, GUCY1A2, GREM1, MACC1, and ABAC13, were upregulated by CHI3L1 but downregulated by KSM (Fig. 6B and Supplemental Table S3). Among these genes, EGFR expression was one of the most prominently regulated genes by CHI3L1 and KSM treatment. Conversely, KSM increased the expression of several genes that were downregulated by CHI3L1, including those with known anti-tumor activities, such as CCL23, NKD2, AZU1, and PLD4 (Supplemental Table S4). In particular, we further confirmed increased mRNA and protein expression of EGFR in CHI3L1-stimulated differentiated M2 macrophages (Fig. 6, C and D). As shown in Fig. 6E, treatment with EGFR inhibitor gefitinib abrogated the CHI3L1-induced macrophage expression of CD163 and CD206, suggesting a role for EGFR in CHI3L1-stimulated macrophage M2 differentiation.

**Figure 6.**
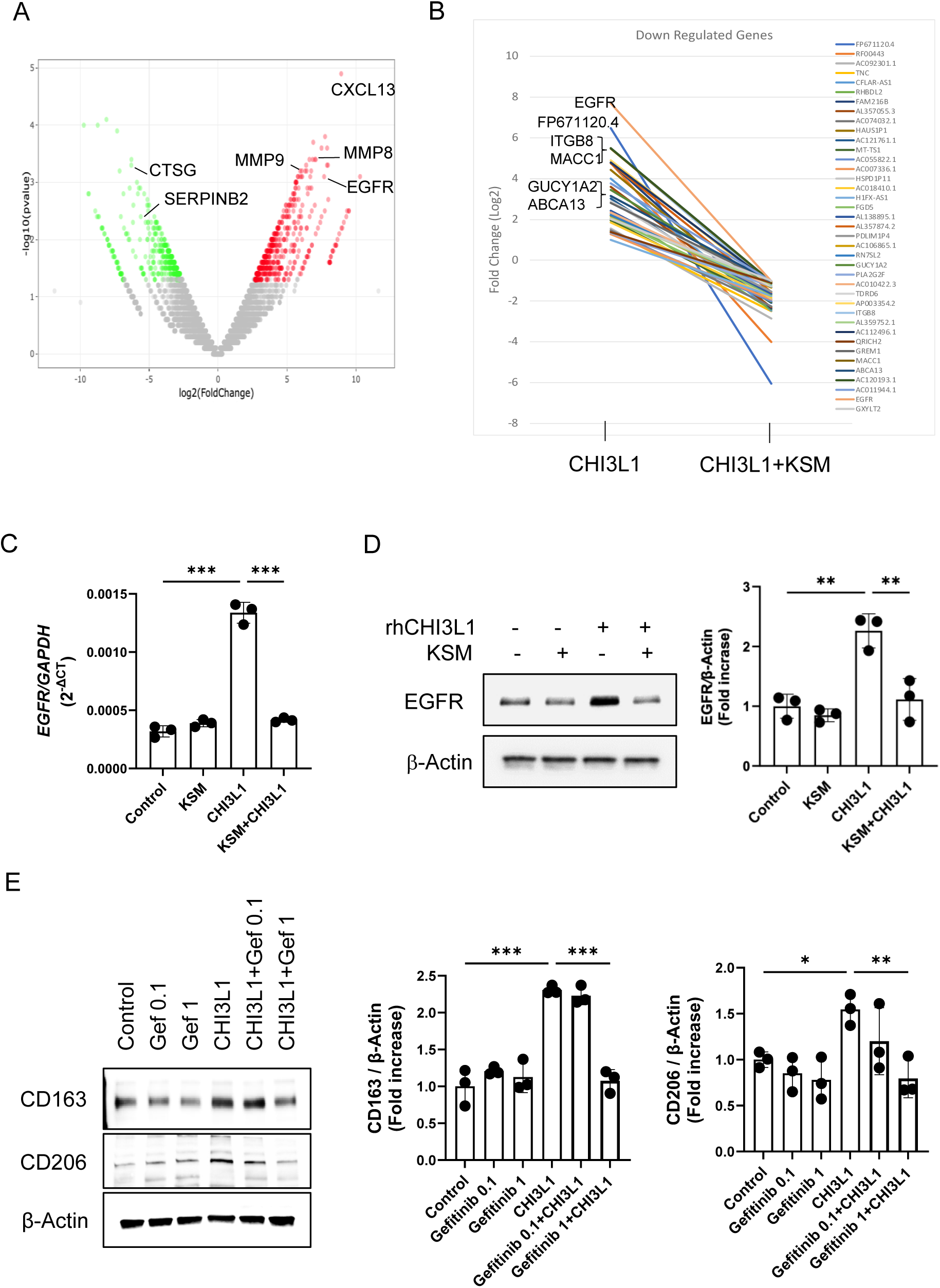
KSM inhibits CHI3L1-stimulated M2 macrophage activation via EGFR. Bulk RNAseq sequencing analysis on the differentiated macrophages from THP-1 cells was used to evaluate mRNA expression in the macrophage differentiation regulated by CHI3L1 and KSM. (A). Volcano plots showing differentially expressed genes regulated by CHI3L1. (B) Representative plots of the top 20 genes that are upregulated (> 2-fold) by CHI3L1 stimulation but downregulated (< 2-fold) by KSM treatment. (C-D) Representative mRNA and protein expression of EGFR in differentiated macrophages with CHI3L1 and KSM treatment detected by real-time qRT-PCR (left panel) and immunoblot evaluations (right panel) in the differentiated macrophages with CHI3L1 and KSM treatments. (E) Effects of EGFR inhibition on the expression of CD163 and CD206 in CHI3L1-stimulated, differentiated macrophages by gefitinib (#3000, Tocris Bioscience, Bristol, UK) treatment (0.1 and 1 μM, 72 hours). Right panel, densitometric quantitation on the blots of CD163 and CD206. The values in panels C-E are mean ± SEM. *p < 0.05, **p < 0.01 ***p < 0.001 (One-Way ANOVA, multiple comparisons).

## Discussion

The tumor microenvironment (TME), composed of diverse immune cells and mediators, plays a crucial role in tumor cell survival, metastasis, and patient prognosis (22–24). Among these, tumor-associated macrophages (TAMs) are predominant innate immune cells in the TME that contribute significantly to immunosuppression, tumor cell evasion, cancer cell proliferation, survival, invasion, and metastasis (25, 26). While exceptions exist, M2 macrophages or M2-like TAMs are generally regarded as pro-tumoral, whereas M1 macrophages or M1-like TAMs are associated with anti-tumor activity (27). Therefore, understanding the mechanisms driving pro-tumoral macrophage activation and identifying regulatory inhibitors is critical for developing interventions against tumor progression. Our findings demonstrated that CHI3L1 drives M2 or M2-like TAM differentiation both *in vivo* and *in vitro*, aligning with its well-established pro-tumoral activities. Moreover, we revealed that Kasugamycin (KSM), an aminoglycoside antibiotic, is an inhibitor of CHI3L1 signaling, bioactivity, and CHI3L1-induced M2 macrophage differentiation. Notably, KSM treatment significantly reduced melanoma lung metastases driven by CHI3L1 overexpression. Together, these results suggest a novel anti-tumor activity of KSM, which likely acts through the inhibition of CHI3L1-stimulated M2-like TAM differentiation. Our studies further identified that the CHI3L1-EGFR axis could be a mechanism underlying CHI3L1-induced M2 or M2-like TAM differentiation, as well as a target of the inhibitory effect of KSM.

Recent work from our laboratory has shown that CHI3L1 is a powerful immune modulator that increases tumor immune tolerance by regulating immune checkpoint molecules in lung macrophages (10, 11). CHI3L1 also promotes alternative macrophage activation, contributing to conditions like pulmonary fibrosis and airway remodeling in asthma (8, 28). However, its direct role in TAM differentiation and its contribution to tumor metastasis and progression had not been well studied. Our current results establish CHI3L1 as a key driver of pro-tumoral M2 macrophage differentiation *in vitro* and in a melanoma lung metastasis model *in vivo*. The role of CHI3L1 in pro-tumoral M2 macrophage activation has also been recognized in other malignancies, including gastric, breast, and lung cancers (29–31). For example, Cohen *et al*. demonstrated that cancer-associated fibroblasts expressing high levels of CHI3L1 enhanced M2 macrophage recruitment and activation, thereby driving breast cancer progression (31). These findings emphasize that CHI3L1-induced pro-tumoral macrophage differentiation represents a promising therapeutic target for disrupting tumor progression.

Despite its translational potential, the development of CHI3L1-targeted therapies for lung and other cancers remains limited. Existing strategies, such as neutralizing anti-CHI3L1 antibodies or gene-specific siRNA approaches, have shown efficacy in preclinical studies (6, 11). While natural compounds like resveratrol have demonstrated anti-CHI3L1 activity (32), their precise molecular mechanisms remain unclear. Targeting putative CHI3L1 receptors or coreceptors, such as IL-13Rα2, TMEM219, CD44, or galectin-3, has also been proposed to inhibit CHI3L1-mediated signaling (19, 33–35). However, these receptors interact with multiple ligands essential for other biological processes, limiting their specificity and clinical use.

Previously, we identified KSM as a chitinase inhibitor through high-throughput screening of chemical and drug libraries (18). The broad-spectrum anti-chitinase activity of KSM has since been validated against various human and microbial chitinases (17). Although CHI3L1 lacks enzymatic activity, its unique chitin-binding domain (CBD), shared with true chitinases, led us to hypothesize that KSM could similarly modulate CHI3L1 bioactivity. It has been shown that chitin-binding domain (CBD) determines the biological function of CHIT1 (36). Evidence of direct physical interactions between KSM and CHIT1 supports this notion, as does the structural similarity of the CBD in CHI3L1 and CHIT1. Recent studies further highlight the critical role of CBD in the biological activity of CHI3L1, particularly in lung and colon cancers (37–39). However, future studies are needed to elucidate the precise KSM binding site on CHI3L1 and how this interaction inhibits its signaling and biological activities.

KSM is a naturally occurring aminoglycoside antibiotic isolated from *Streptomyces kasugaensis* (40). Unlike other aminoglycosides, KSM has a unique carbohydrate structure comprising two sugars, a D-chiro-inositol moiety and a kasugamine moiety (2,4-diamino-2,3,4,6-tetradeoxy-D-arabino-hexose) with a carboxy-imino-methyl group (41). These distinctive features raise the possibility of a direct interaction between KSM and the carbohydrate-binding domains of CHI3L1. While originally developed as an antibiotic effective against a broad range of microorganisms, KSM is currently used as a fungicide in organic agriculture due to its low toxicity in plants, humans, fish, and animals (42, 43). Its past use in treating respiratory and urinary tract infections (44, 45) highlights its potential clinical utility. Developing KSM as an anti-cancer drug offers significant advantages, including drug accessibility for patients with lung cancer. However, as an aminoglycoside, KSM’s impact on the microbiome warrants careful consideration. Future efforts should focus on developing KSM derivatives with minimized antibiotic activity while retaining their anti-chitinase and anti-tumor properties.

Transcriptomic analysis of CHI3L1-stimulated genes in M2 macrophage differentiation provided further insight into the molecular basis of KSM’s anti-tumoral activity. CHI3L1 upregulated several pro-tumoral genes while downregulating genes with anti-tumor activity. Among these, EGFR was particularly noteworthy, as its expression was significantly stimulated by CHI3L1 and suppressed by KSM. EGFR is a well-established driver of tumor progression in lung and other cancers (46–48), and its signaling in macrophage activation has been implicated in infection and tumor development (46, 49). Recently, our lab reported a feed-forward relationship between CHI3L1 and EGFR expression has been reported in normal or EGFR mutant lung epithelial cells (50). However, whether a similar CHI3L1-EGFR axis exists in immune cells such as macrophages has remained unexplored. Our present study is the first to demonstrate that CHI3L1 upregulates EGFR expression in macrophages, leading to M2-like TAM differentiation. This novel axis represents a previously unrecognized pathway by which CHI3L1 promotes tumor immune evasion, and KSM’s ability to counteract this signaling suggests a promising therapeutic strategy. Interestingly, KSM has also demonstrated antiviral immune activity by enhancing type I interferon responses (51), suggesting that it may indirectly augment macrophage-mediated anti-tumor immunity. Taken together, these studies suggest that the anti-tumor effect of KSM was potentially mediated through interruption of the CHI3L1-EGFR axis that plays an essential role in macrophage polarization and/or differentiation. The effects of KSM on macrophage functions, such as phagocytosis and interactions with cytotoxic T cells, remain to be determined.

In summary, our findings demonstrate that KSM inhibits CHI3L1-mediated signaling and oncogenic M2 macrophage differentiation, highlighting its potential to counteract CHI3L1’s protumor activities. These results position KSM as a promising small molecule for development as a first-in-class chitinase-inhibitor-based anti-cancer agent, particularly for cancers where CHI3L1 plays a critical role.

## Materials and Methods

### Genetically Modified Mice

Lung-specific CHI3L1-overexpressing transgenic mice (CHI3L1 Tg; *Chil1* Tg), in which CHI3L1 is targeted to the lung using the CC10 promoter, were generated and characterized in our laboratory as previously described (8, 33). Experiments were conducted using 8–10-week-old wild-type (WT) and *Chil1* Tg mice. All protocols were reviewed and approved by the Institutional Animal Care and Use Committee (IACUC) at Brown University.

### Melanoma Lung Metastasis and KSM Treatment

The B16-F10 mouse melanoma cell line (#CRL-6475, ATCC, Manassas, VA, USA) was cultured in Dulbecco’s Modified Eagle Medium (DMEM) supplemented with 10% fetal bovine serum (FBS) and 1% penicillin-streptomycin. Cells were collected upon reaching 80% confluence, adjusted to a concentration of 1×10^6^ cells/mL, and injected into mice via the lateral tail vein (2×10^5^ cells/mouse in 200 μL DMEM; low-dose challenge, 0.5×10^5^ cells/mouse) (11). Intraperitoneal KSM (#K4013, MilliporeSigma, Burlington, MA, USA) injections at specified doses were initiated on Day 2 after the tumor cell challenge and administered every other day for two weeks. Lung metastases were quantified by counting melanoma colonies (black nodules) on the pleural surface as described previously (7, 9).

### FACS analysis

Single-cell suspensions from whole mouse lungs were prepared using the Lung Dissociation Kit (Miltenyi Biotec Inc., Auburn, CA, USA) per the manufacturer’s instructions. Cells were stained with fluorescently labeled antibodies directed against CD45 (30-F11, #103126, Thermo Fisher Scientific, Waltham, MA, USA), PD-L1 (MIH5, #12-5982-8, Thermo Fisher Scientific), CD11b (M1/70, #101210, BioLegend, San Diego, CA, USA), CD206 (MR5D3, #MA5-16870, Thermo Fisher Scientific), CD163 (TNKUPJ, #17-1631-82, Thermo Fisher Scientific). Flow cytometry data were collected using the BD FACSAria IIIu and analyzed with FlowJo (v10) software. The gating strategy of the macrophage is illustrated in Supplemental Fig. S5.

### *In Vitro* Macrophage Differentiation Using CHI3L1 And Tumor Cell Conditioned Media

Human monocytic THP-1 cells were differentiated into TAMs using recombinant (r) CHI3L1 along with tumor cell-conditioned media (from A549 cancer cells), following established procedures with modification (52). In this protocol, we replaced the traditional IL-4/IL-10 cytokines with CHI3L1. KSM effects were evaluated by adding 250 ng/mL KSM to the TAM differentiation medium. Differences in TAM differentiation (measured by CD206, CD163, and PD-L1 co-expression) were assessed by real-time qRT-PCR, FACS, and immunoblot evaluations. The expression of other markers of macrophage activation, such as CX3CR1, CD36, and NOS2, was also evaluated by real-time RT-PCR. The specific role of CHI3L1 in macrophage differentiation was also investigated using CHI3L1-neutralizing monoclonal antibody (FRG antibody).

### RNA Extraction and Semi-Quantitative Real-Time qPCR

Total RNA was isolated using TRIzol reagent (ThermoFisher Scientific) followed by RNA purification with the RNeasy Mini Kit (Qiagen, Germantown, MD, USA) per the manufacturer’s instructions. Reverse transcription and real-time PCR were performed as previously described (8, 53). Ct values for target genes were normalized to internal housekeeping genes GAPDH. Primer sequences are listed in Supplemental Table S5.

### Bulk RNA Sequencing and Data Analysis

Macrophages differentiated from THP-1 cells under various treatments were subjected to bulk RNA sequencing. Total RNA isolated from pooled duplicated samples were subjected to the evaluation. The differential transcriptomic gene expression profiling and data analysis was conducted to identify genes regulated by CHI3L1 and KSM using illuminar’s next generation sequencing that detect coding and long non-coding RNAs (Standard RNA-Seq Service provided by Genewiz/Azenta Life Science, Waltham, MA, USA).

### Western Blotting (Immunoblotting)

Protein lysates (25 µg) from lung tissues or cells were subjected to SDS-PAGE, transferred to membranes, and immunoblotted with primary antibodies against CD163 (#PA5-109327, Thermo Fisher Scientific), CD206 (E6T5J, #24595S, Cell Signaling Technology, Danvers, MA, USA), phosphorylated AKT (pAkt) (193H12, #4058S, Cell Signaling Technology), and total Akt (11E7, #4685S, Cell Signaling Technology), EGFR (D38B1, #4267S, Cell Signaling Technology), phosphorylated Erk (pErk) (#9101S, Cell Signaling Technology), total Erk (#9102S, Cell Signaling Technology), and β-actin (C4, #sc-47778 HRP, Santa Cruz Biotechnology, Dallas, TX, USA). HRP-conjugated anti-rabbit IgG secondary antibody (#7074S, Cell Signaling Technology) was used for detection. Immunoblotting was performed following standard protocols as described previously (54).

### Measurement of Oxidative Stress-Induced Apoptosis

Cells were treated with hydrogen peroxide (1 mM, 24 hours, #216763, Sigma-Aldrich, St. Louis, MO, USA) to induce oxidative stress. Apoptotic cells were labeled with Annexin V and propidium iodide (#556547, BD Biosciences, San Jose, CA, USA), followed by flow cytometry analysis.

### Double-Label Fluorescent Immunohistochemistry

Formalin-fixed, paraffin-embedded (FFPE) lung tissue blocks were sectioned into 5 µm-thick slices and mounted onto glass slides. Sections were deparaffinized, rehydrated, and subjected to heat-induced epitope retrieval using a steamer and antigen unmasking solution (Abcam, citrate buffer, pH 6.0) for 30 minutes. Blocking was performed with serum-free protein blocking solution (Dako, Agilent Technologies, Santa Clara, CA, USA) for 60 minutes at room temperature. Slides were incubated overnight at 4°C with primary antibodies against CD163 (EPR19518, #ab182422, Abcam, Cambridge, UK), CD206 (#AF2535, R&D Systems, Minneapolis, MN, USA). After washing, fluorescent-labeled secondary antibodies of Donkey anti-Rabbit IgG (H+L) Alexa Fluor™ 488 (#A-21206, Thermo Fisher Scientific) or Donkey anti-Goat IgG (H+L) Alexa Fluor™ 594 (#A-11058, Thermo Fisher Scientific) were applied for 1 hour at room temperature. Sections were counterstained with DAPI and mounted with coverslips.

### Quantification and Statistical Analysis

All statistical analyses were conducted using GraphPad Prism software. Comparisons between groups were made using two-tailed Student’s t-tests. One-way ANOVA followed by appropriate post-hoc tests was performed for multiple group comparisons. Data are presented as mean ± SEM, and statistical significance was defined as p < 0.05.

## Data availability statement

The bulk RNA sequencing data supporting the findings of this study are available in the Supplementary Materials.

